# The novel HP1a interactor Clump/CG30403 safeguards transposon silencing and reproductive resilience in the *Drosophila* ovary

**DOI:** 10.64898/2026.07.15.738798

**Authors:** Kun Wu, Richard Chang, Andrew Garcia, Jian Fang, Jikui Song, Maria Ninova

**Affiliations:** Department of Biochemistry, University of California Riverside 3401 Watkins Drive, Boyce Hall 1415A Riverside, CA 92521, USA

## Abstract

Heterochromatin and its core effectors from the Heterochromatin Protein 1 (HP1) family are essential for epigenetic silencing of repetitive regions and genome integrity from yeast to humans. However, how HP1s and associated factors regulate heterochromatin properties *in vivo* to sustain silencing across development, aging, and environmental factors, remains incompletely understood. Here, we identify Clump/CG30403 — a previously uncharacterized MADF-BESS domain protein — as a novel heterochromatin factor required for robust transposon silencing during *Drosophila* oogenesis and sustained fertility with age and temperature stress. Clump/CG30403 interacts with the main HP1-family silencing effector Su(var)205/HP1a through a noncanonical binding motif within its large intrinsically disordered region. Notably, in the absence of Clump/CG30403, HP1a mobility and silencing capacity are compromised despite largely unperturbed genomic distribution, showing that HP1a presence alone is insufficient for repression. We also show that Clump/CG30403 uniquely accumulates at its own promoter to self-repress and prevent ectopic aggregation, revealing a feedback mechanism to constrain protein dosage and phase behavior. Overall, we propose that Clump/CG30403 is a HP1 corepressor that acts as a tightly calibrated safeguard of the heterochromatin environment properties to ensure stable silencing, genome integrity, and persistent reproductive function.

## Introduction

Heterochromatin, the highly compacted form of chromatin enriched at repetitive genomic regions and developmentally silenced genes, plays critical structural and regulatory roles in safeguarding genome integrity and transcriptome fidelity, including restricting the activity of transposable elements (TEs) and lineage-inappropriate genes, and supporting proper chromosome structure and segregation(Grewal and Jia 2007; Elgin and Reuter 2013; Allshire and Madhani 2018; Grewal 2023). From yeast to human, the bulk of constitutive heterochromatin — including telomeric and pericentromeric repeats, interspersed transposable elements (TEs), as well as a subset of developmentally silenced regions — is regulated by histone 3 lysine 9 trimethylation (H3K9me3) and the Heterochromatin Protein 1 (HP1) family(Grewal and Jia 2007; Allshire and Madhani 2018; Janssen et al. 2018). First identified as Su(var)205 in pioneering studies in *Drosophila*, HP1 proteins are now recognized as highly conserved readers of methylated H3K9 and central effectors of nucleosome compaction, promoting a condensed environment classically viewed as restrictive to transcription initiation(James and Elgin 1986; Bannister et al. 2001; Lachner et al. 2001; Nakayama et al. 2001; Jacobs and Khorasanizadeh 2002; Machida et al. 2018; Sanulli et al. 2019; Lou et al. 2024). Furthermore, HP1 can interact with H3K9 writers in a feedforward process that ensures heterochromatin maintenance and spreading, and HP1 proteins act as potent transcriptional silencers when recruited to chromatin(Schotta et al. 2002; Li et al. 2003; Hathaway et al. 2012; Al-Sady et al. 2013). Nevertheless, H3K9me3/HP1-marked heterochromatin is neither structurally static nor uniformly repressive. Biophysically, HP1 proteins can phase separate, and heterochromatin domains display liquid-liquid phase separation (LLPS) behavior with a mobile HP1a fraction(Larson et al. 2017; Strom et al. 2017; Sanulli et al. 2019). Functionally, HP1 occupancy does not always equate absolute transcriptional repression, and some heterochromatic regions encode functional products(Wakimoto and Hearn 1990; Riddle et al. 2011; Riddle et al. 2012). In the *Drosophila*germline, for example, non-canonical transcription from specialized heterochromatic regions, co-occupied by the principal heterochromatin-associated HP1 protein Su(var)205/HP1a and its paralog Rhino/HP1d, produce piRNAs that direct both the post-transcriptional and co-transcriptional silencing of TEs(Brennecke et al. 2007; Klattenhoff et al. 2009; Sienski et al. 2012; Le Thomas et al. 2013; Rozhkov et al. 2013; Mohn et al. 2014; Teo et al. 2018). Together, these biophysical and functional nuances raise the important question of how different HP1 proteins engage with other heterochromatin proteins to achieve appropriate, locus-specific assembly and dynamics. Here, we characterize CG30403 — a protein with unknown function but previously identified as both a physical interactor and regulatory target of Su(var)205/HP1a in *Drosophila melanogaster(Alekseyenko et al. 2014; Ninova et al. 2020)*. We named this protein Clump for *chromatin-linked uncharacterized MADF-BESS protein* and for its aggregation behavior described later in this study. Clump/CG30403 belongs to the MADF-BESS family which shares a conserved architecture consisting of an N-terminal MADF domain, related to Myb/SANT-like DNA-binding modules, and a C-terminal BESS domain that mediates protein-protein interactions, connected by variable unstructured linker regions(Shukla et al. 2014). Consistent with this architecture, characterized members of this family include transcription factors and chromatin regulators, and their expansion to over 16 paralogs in the Drosophilid lineage suggests functional specialization for distinct genomic targets and regulatory roles(Bhaskar and Courey 2002; Shukla et al. 2014; Zinshteyn and Barbash 2022; Chavan et al. 2023).

We show that Clump is specifically expressed in the *Drosophila* ovary and interacts with the main heterochromatin effector HP1a via its large intrinsically disordered region (IDR). Loss of Clump causes variegated activation of selected TEs and reduced female fertility, with these effects worsening with aging and temperature stress, consistent with cumulative chromatin instability. Mechanistically, we show that Clump is recruited to heterochromatin by HP1a and is required for its mobility and efficient silencing. Defects in both canonical and non-canonical transcription upon Clump loss point to underlying chromatin perturbations that converge on TE derepression, despite buffering by a largely intact piRNA pathway, thereby positioning Clump as a HP1a corepressor acting downstream of piRNA-mediated targeting to reinforce heterochromatin silencing. Interestingly, Clump is highly prone to aggregation — a property conferred by its IDR-BESS region — and, unlike HP1a, Clump has limited mobility *in vivo*. At physiological levels, Clump foci are exclusively nuclear and co-localized with HP1a, but its overexpression results in aberrant cytoplasmic aggregates. We demonstrate that in addition to genome-wide localization to HP1a-occupied heterochromatin, Clump can bind its own promoter sequence via the DNA-binding MADF domain and self-repress, thereby establishing a negative-feedback loop that controls its abundance. Together, these findings position Clump within a multilayered regulatory network that integrates chromatin targeting, protein dynamics, and autoregulatory feedback to ensure robust and precisely tuned heterochromatin regulation in the germline necessary to safeguard female fertility.

## Results

### Clump interacts with HP1a via its IDR and localizes across global heterochromatin in the *Drosophila* ovary

The *clump/cg30403* locus encodes a 33.4 kDa protein where the structured MADF and BESS domains are connected through a long intrinsically disordered region (Figure 1A). To further characterize this protein, we knocked in a C-terminal eGFP-3xFLAG tag by CRISPR/Cas9-mediated homology-directed repair (HDR) and isolated homozygous *clump[KI.eGFP::3xFLAG]* strains alongside wild-type sibling control strains, herein referred to as Clump-GF and WT, respectively (Figure 1B). Consistent with available gene expression atlas data(Ozturk-Colak et al. 2024), Western Blot (WB) analysis showed restricted expression pattern to ovaries and early embryos, indicating a role in oogenesis (Figure 1C). At the subcellular level, Clump-GF displays a clear nuclear localization across the germline throughout oogenesis, starting in the germarium, with pronounced nuclear foci in nurse cells and a single bright focus in the oocyte; a weaker but distinct signal is also present in the somatic follicle cells (Figure 1D).

**Figure 1.**
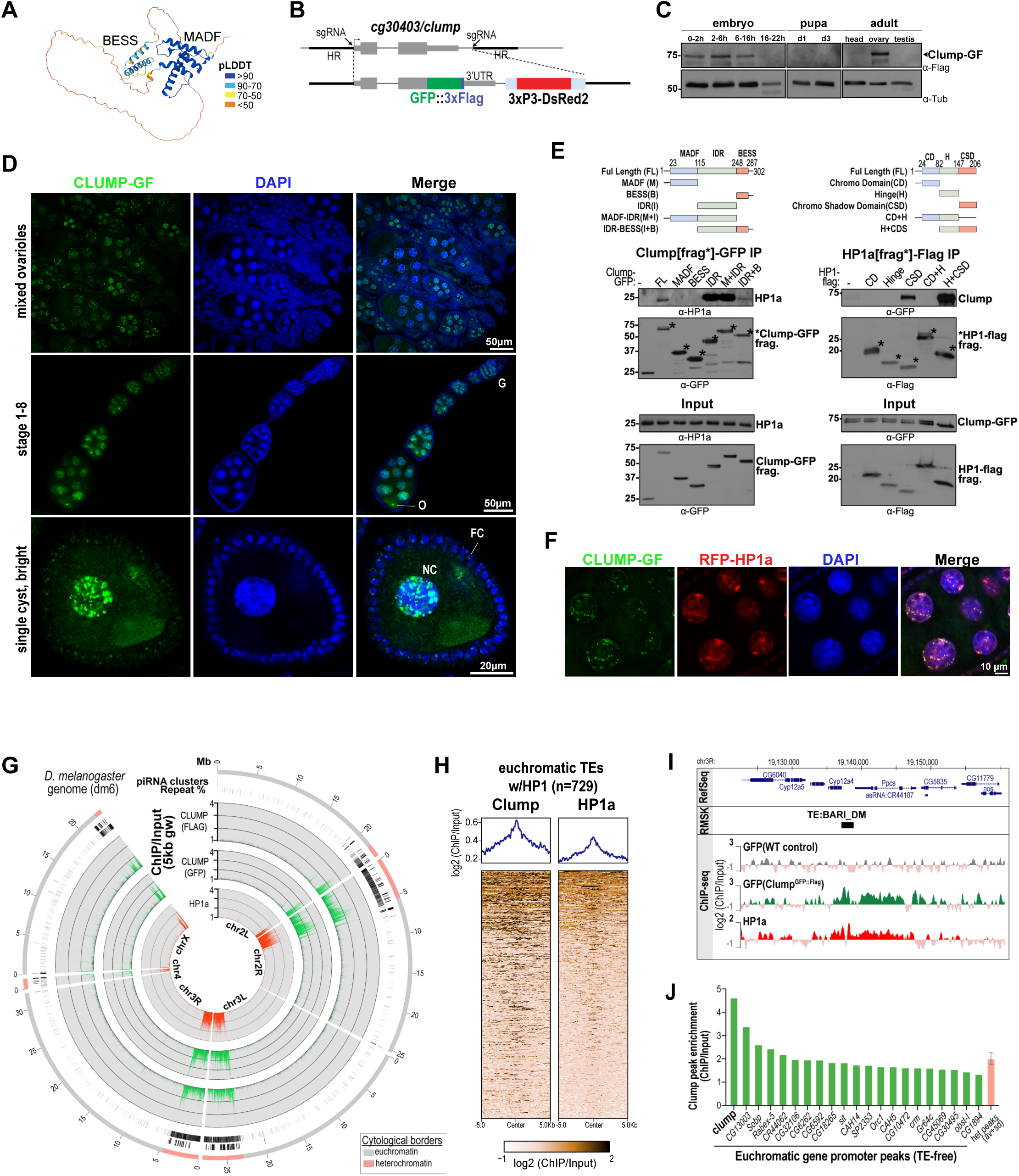
Clump colocalizes with HP1a across heterochromatin and TE insertions in the *Drosophila* ovary. **(A)** AlphaFold3-predicted model of Clump protein, color coded according to the pLDDT score; the BESS and MADF domains of Clump are labelled. **(B)** Schematic of the CRISPR/Cas9 strategy used to generate the endogenous Clump-GF allele. **(C)** WB analysis of Clump expression in indicated tissues and stages. **(D)** Confocal images of Clump-GF (green) in the ovary at different magnifications. NC=nurse cells, FC=follicle cells, O=oocyte; G=germarium. **(E)** WB analysis of reciprocal Clump and HP1 co-immunoprecipitation experiments. Constructs of indicated tagged fragments of Clump and HP1a fragments were overexpressed in S2 cells. **(F)** Confocal images of nurse cell nuclei showing nuclear co-localization of RFP-HP1a and Clump-GF. **(G)** Circle plot of ChIP-seq data showing genome-wide distribution of HP1a and Clump-GF ChIP to Input signal. The outer ideogram represents the chromosome arms of Drosophila melanogaster (dm6), with cytologically defined euchromatic and heterochromatic regions marked. Adjacent inner tracks indicate piRNA clusters and repeat density. Inner tracks show ChIP-seq enrichment from experiments using anti-FLAG and anti-GFP antibodies to detect Clump (green) and anti-HP1a antibodies (red). ChIP-seq controls for anti-FLAG and anti-GFP signals using wild-type control ovaries that do not express Clump-GF shown in grey. Signals are displayed as RPKM-normalized ChIP/Input ratios in 5-kb genomic windows. **(H)** Heatmap showing Clump and HP1a ChIP-Seq enrichment across genomic regions flanking euchromatic transposable element insertions. The heatmap is centered on 729 reference TE insertions located in euchromatin overlapped with HP1a MACS2 peaks. Each rows represent individual TE insertion and are ordered by decreasing Clump and HP1a ChIP-seq enrichment. **(I)** UCSC Genome Browser screenshot of a locus harboring reference TE insertion. Tracks show log2-transformed ChIP/Input enrichment profiles generated from uniquely mapped reads from ovarian ChIP-seq samples. Clump-GF signal was detected by GFP ChIP-seq, using wild-type ovaries lacking Clump-GF as the GFP ChIP-seq negative control. HP1a ChIP-seq signal is shown for comparison. Images are representative of 2-3 biological replicates. **(J)** ChIP-seq enrichment of Clump at peaks annotated to gene promoters and at heterochromatin peaks. Heterochromatic peak enrichment was calculated as the mean signal across MACS2-identified heterochromatic Clump peaks, with error bars indicating standard deviation.

Clump was previously identified as an HP1a interactor(Alekseyenko et al. 2014). Consistent with this, Clump-GF co-purifies with HP1a in ovarian lysates (Figure S1A). Through series of reciprocal coimmunoprecipitation experiments using different truncated fragments of the two proteins in *Drosophila* S2 cells, we further demonstrate that the interaction between Clump and HP1a is sufficiently mediated by Clump’s IDR region and HP1a’s chromoshadow (CSD) domain (Figure 1E). It is well-established that HP1 CSDs form a symmetrical homodimer that creates a hydrophobic groove to harbor the consensus pentapeptide motif PxVxL(Smothers and Henikoff 2000; Thiru et al. 2004; Huang et al. 2006; Mendez et al. 2013; Canzio et al. 2014; Liu et al. 2017; Shao et al. 2024). This understanding was recently expanded revealing a more degenerate and extended array of CSD binding sequences containing hydrophobic and basic residues, termed HP1a Access Codes (HACs)(Colmenares et al. 2025). Despite the largely unstructured nature of Clump’s IDR, AlphaFold3(Abramson et al. 2024) modeling predicts an interaction between the HP1a CSD and residues 129-133 of Clump, residing in a conserved segment of the IDR (Figure S1B,C). Notably, this segment occupies the canonical protein-interaction sites at the homodimeric interface of the HP1 CSD, where it pairs with the C-terminal β-strands of each CSD monomer into a three-stranded β-sheet, resembling previously characterized structures(Smothers and Henikoff 2000; Thiru et al. 2004; Huang et al. 2006; Mendez et al. 2013; Canzio et al. 2014; Liu et al. 2017; Shao et al. 2024)(Figure S1B,C). The L129-I131-L133 motif of the Clump IDR engages in non-polar contact with the surface groove of the CSD, in a manner similar to the canonical contact between the PxVxL motif and the CSD dimer. Our sequence analysis further revealed that the CSD interacting motif of the Clump IDR is highly conserved among *D. melanogaster* and most other drosophilids(Figure S1B). One exception is the *obscura* group where homologs carry an I>V substitution, highlighting a degree of plasticity, consistent with the findings that the central I and V are interchangeable(Colmenares et al. 2025). Collectively, the experimental data and structural similarity to established HP1a binding interfaces identify Clump as a *bona fide* HP1a interactor.

Building on these observations, we next examined the spatial relationship between Clump and HP1a *in vivo* throughout ovary development. Confocal imaging reveals largely overlapping signals of the two proteins, indicating that Clump globally co-localizes at heterochromatin alongside HP1a (Figure 1F). To resolve Clump’s chromatin occupancy and its relationship to HP1a at higher resolution, we performed ChIP-seq on WT and Clump-GF ovaries using FLAG and GFP epitopes in independent experiments, alongside HP1a ChIP-seq. Results yielded reproducible patterns of Clump genome-wide localization closely mirroring that of HP1a. Specifically, abundant Clump and HP1a signals are present at the repeat-rich cytological heterochromatin regions(Riddle et al. 2011), including chromosome 4 and the pericentromeric regions of chromosomes 2, 3, and X (Figure 1G); interestingly, Clump displays a slight enrichment bias to the pericentromeric heterochromatin/euchromatin boundaries of the 2nd and 3rd chromosomes, compared to the more uniformly distributed HP1a. While telomeric regions are not in the reference assembly and hence are inaccessible for ChIP-seq analysis, co-staining with the capping factor HipHop confirmed Clump also localizes to telomeres (Figure S1D). Additionally, Clump is present on dual-strand piRNA clusters — large heterochromatic islands bound by HP1a and its paralog Rhino, that are enriched in TE fragments and act as sources of precursor piRNA transcripts — but absent from euchromatic, HP1a-independent piRNA clusters such as *tj* (Figure S1E). Further, Clump occupancy follows HP1a at interspersed TEs within the euchromatic chromosomal arms (Figure 1H,I). Collectively, these findings establish Clump as a ubiquitous partner of HP1a that co-localizes at HP1a-controlled sites in the *Drosophila* ovary.

Finally, we identified a limited set of 45 Clump peaks in euchromatic regions lacking TE insertions within 5 kb, 22 of which reside in gene promoter regions (Figure 1J, S1F). While most of these peaks have relatively modest signals, this subset contains the single most significant Clump peak in the entire dataset — exceeding the typical genome-wide Clump-GF ChIP signal more than 2-fold. Strikingly, this specific peak maps to the *clump* gene promoter, pointing to an auto-regulatory mechanism further detailed in the last section (Figure 1J, also see Figures 5A,C).

### Clump loss causes reduced fertility and variegated TE activation

To gain an insight into the function of Clump, we sought to determine the consequences of its loss in mutant and knockdown strains. Unexpectedly, the available RNAi lines and several deficiency stocks for this gene appeared to have lost respective transgenes and deletions (data not shown). We therefore generated a new *clump* null mutant strain *clump[Δ::dsRed]* by precise excision and exchange of the entire gene with a dsRed cassette via CRISPR-Cas9 HDR (Figure 2A). Of note, although RefSeq annotations suggest that *clump/cg30403* is nested in an intron of a long *hmgD* isoform, close inspection showed that this isoform is likely a mis-annotation, and importantly, the *clump* deletion does not affect the expression of neighboring gene isoforms (Figure S2A). We also generated two strains encoding UASp-driven small hairpin RNAs for *clump* conditional knockdown in the germline, the stronger of which (shClump-1) was used in downstream experiments (Figure S2B).

**Figure 2.**
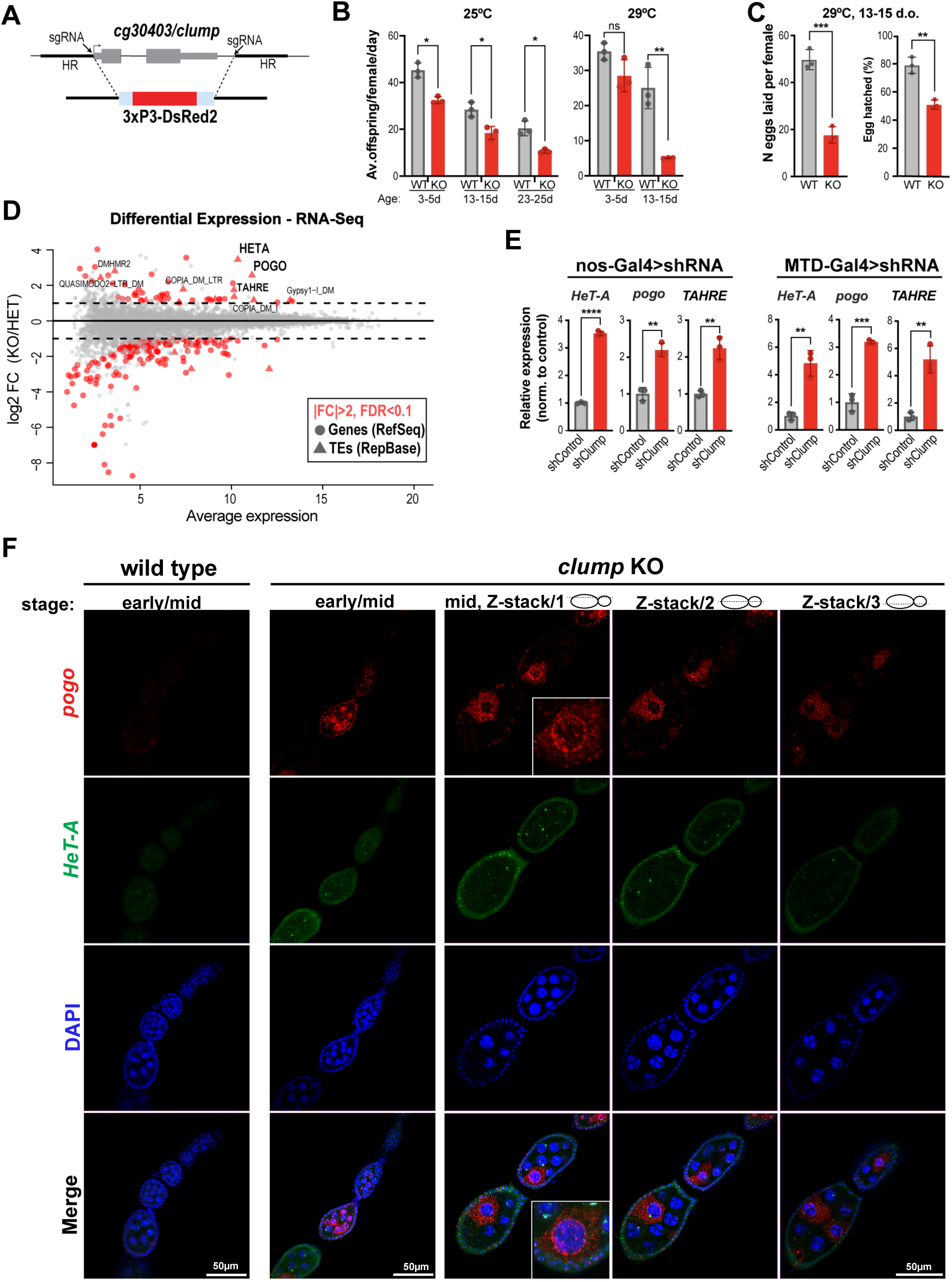
Loss of Clump impairs fertility and causes transposon upregulation. **(A)** A schematic diagram illustrating the CRISPR-Cas9-mediated knockout of *clump* via HDR repair. **(B)** Female fertility assays for *clump* knockout flies. Indicated female genotypes were crossed to wild-type (yw) males, and fertility was assessed by quantifying the number of adult F1 progeny produced per female per day. Fertility was measured at the indicated female ages and temperatures. Wild-type (yw) females served as controls. Data are shown as mean ± s.d. from biological replicates. Statistical significance was determined by two-tailed Student’s t-test; ns, not significant; *P < 0.05, **P < 0.01, ***P < 0.001. **(C)** Egg-laying and egg-hatching assays for control and *clump* knockout females under the 29°C condition. **(D)** Differential expression analysis of genes and RepBase consensus transposable elements in poly(A)-selected RNA-seq libraries; n=3. **(E)** RT-qPCR analysis of transposon and histone genes in the control (shWhite) and germline knockdown conditions, expression levels were first normalized to *rp49* for each biological replicate. The normalized values were then divided by the mean normalized expression of the control group, which was set to 1. Individual points represent biological replicates, and error bars represent mean and SD. *: p<0.05, **: p<0.01, ***: p<0.001(Student’s t-test). **(F)** RNA *in situ* HCR against *HeT-A* and *pogo* in control and *clump* knockout ovaries. DNA was stained with DAPI (blue).

Overall, *clump* null mutants are viable and sufficiently fertile to survive under standard rearing conditions, and mutant ovaries and embryos appear morphologically normal. However, *clump* loss results in significantly reduced female fertility compared to wild-type siblings at standard temperature (25°C), which progresses to near complete sterility with age and when flies are reared under temperature stress conditions, i.e., 29°C (Figure 2B). The fertility defect manifests as both reduced egg laying and egg hatching (Figure 2C), without an obvious stage-specific arrest. Collectively, these findings indicate that Clump is not strictly essential for a specific developmental checkpoint, but its loss may lead to stochastic defects throughout oogenesis, ultimately compromising overall fertility.

To elucidate the consequences of *clump* loss on the molecular level, we next examined the differences in gene expression between the *clump* null mutants (“KO”) and heterozygous sibling controls (“HET”, *clump[Δ::dsRed]/CyO*) using RNA sequencing (RNA-seq) after polyA selection. Consistent with the overall absence of Clump from gene-rich regions, differential expression analysis demonstrated a limited effect on gene expression, with ∼25 and ∼80 genes being more than 2-fold up– or down-regulated, respectively (Figure 2D). However, *clump* loss is associated with the upregulation of several TE families, with the largest changes in *HeT-A,* the main component of *Drosophila* telomeres, and the less well-characterized DNA transposon *pogo.* (Figure 2D, S2C). *TAHRE* — the second of the three telomeric TEs, and several LTR TEs including *Copia*, *Stalker, Gypsy,* are also among the significantly upregulated set (Figure 2D, S2C). This phenotype is reproducible when Clump is conditionally depleted from the germline using *clump* shRNA under the control of the Maternal Triple GAL4 (MTD-GAL4) or the nanos-GAL4 drivers, indicating that the observed results are independent from the genetic background (Figure 2E). Notably, the prominent telomeric TE activation in KO ovaries phenocopies HP1a loss in terms of telomeric TEs being the most highly sensitive elements to this loss(Teo et al. 2018), supporting a model where Clump functions in concert with HP1a.

We further assessed the spatial activation of the most upregulated TEs *HeT-A* and *pogo* in *clump* KO ovaries by Hybridization chain reaction (HCR). Transcripts of both TEs are barely detectable in WT ovaries, but show clearly increased signals in the KO with distinct patterns: *HeT-A* displays prominent puncta in the nuclei of germline and somatic cells, while *pogo* is prominent in the cytoplasm of germarium and nurse cells, where it distinctly concentrates at the perinuclear region, which may reflect transcripts captured for processing by the ping-pong pathway (Figure 2F). Notably, *pogo* displays a strongly variegated expression pattern with only a subset of neighboring cells in each cyst having strong HCR signal, indicative of a stochastic loss of epigenetic silencing that may be clonally inherited. Together, these spatial profiles suggest that Clump is a heterochromatin regulator required to confer robust silencing of repetitive elements in distinct genomic regions throughout oogenesis.

### Clump loss results in local changes of piRNA populations without global disruption of the piRNA pathway

The piRNA pathway provides the primary mechanism that identifies TE for transcriptional and post-transcriptional silencing in the *Drosophila* ovary. Given the localization of Clump on piRNA clusters and the TE upregulation phenotype in *clump* KO, we investigated the effect of *clump* loss on the piRNA complement by small RNA sequencing. Overall, *clump* loss does not cause a major shift in small RNA size distributions, nucleotide biases, or ping-pong signatures (Figure S3A). Uniquely mapping piRNA reads corresponding to most major piRNA clusters were also largely unperturbed (Figure S3B), suggesting that Clump is not an essential piRNA biogenesis factor.

However, analysis of unique sequences flanking euchromatic TE insertions from the dm6 reference revealed site-specific increases in mapped piRNA in the *clump* KO (Figure 3A,B). Rather than a direct effect, this may indicate compensatory engagement of the piRNA pathway actively attempting to counter TE derepression when Clump is depleted. We also detected local changes in some piRNA clusters, such as a ∼40% decrease in unique piRNA reads corresponding to the 80EF cluster, which is the most highly occupied piRNA cluster by Clump (Figure S1C, S3B, S3C). Furthermore, we observed an increase in spliced transcripts from within the 42AB cluster, indicative of a shift in the balance of canonical versus non-canonical transcription in this region — a hallmark of disrupted cluster epigenetic environment(Zhang et al. 2014; Chang et al. 2019) (Figure S3D). Interestingly, while locus-specific analysis of piRNA profiles is limited by their short size and repetitive nature, TE consensus-level analysis revealed a highly specific, substantial loss of sense and antisense piRNAs corresponding to *pogo* (Figure 3B). While we cannot pinpoint the precise genomic source of the majority of anti-*pogo* piRNAs due to the sequence repetitiveness and incomplete genome assembly, their selective loss implies that the primary *pogo* piRNA source loci reside within chromatin domains highly sensitive to Clump loss.

**Figure 3.**
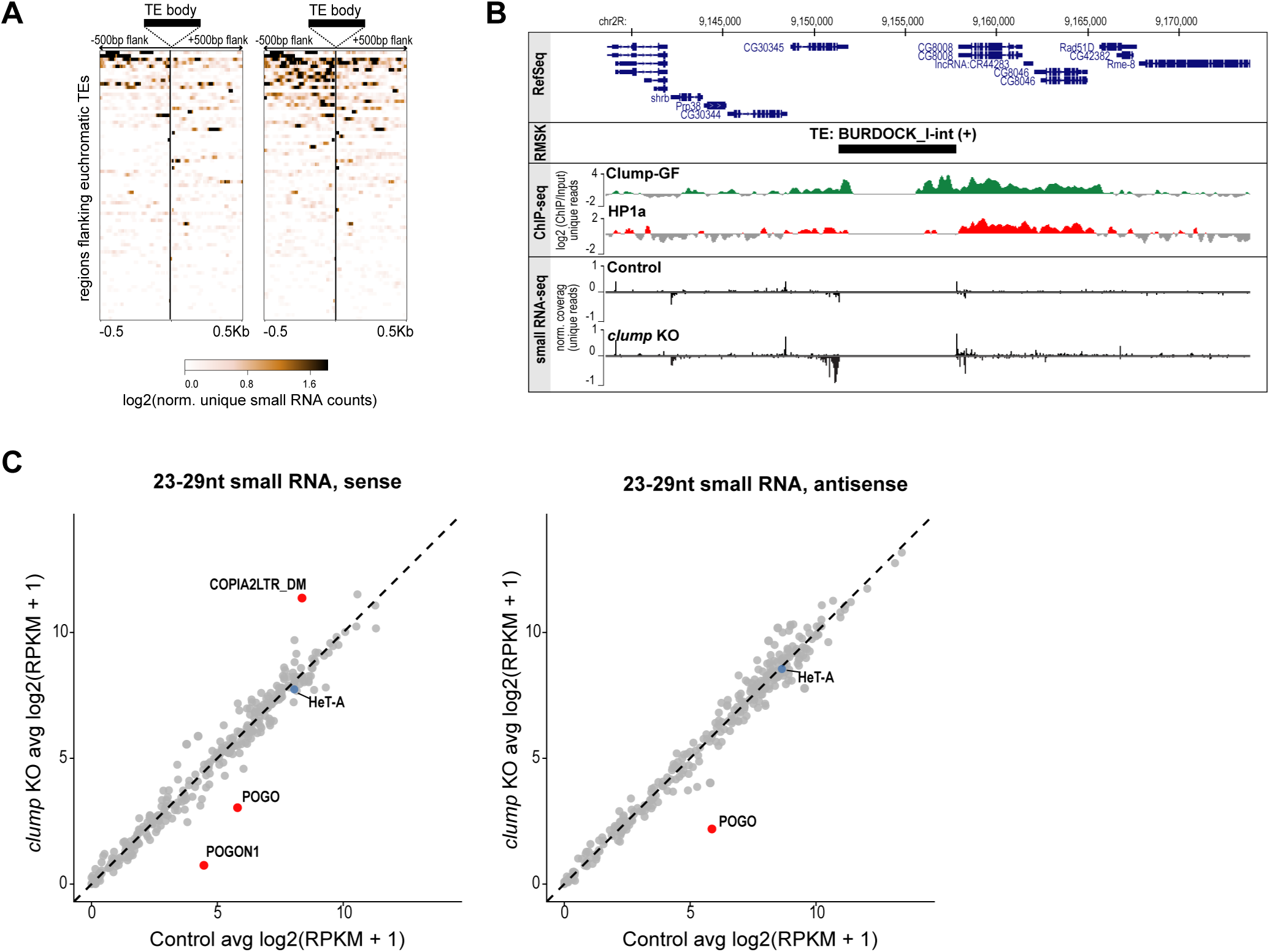
Local effects of Clump depletion on piRNA biogenesis. **(A)** Heatmaps showing Clump and HP1a small RNA sequencing signal in indicated samples 500 bp genomic regions flanking euchromatic TE insertions (without signal in the TE body) in *Drosophila* ovaries. Values are average of 3 biological replicates **(B)** Genome browser view of a representative TE insertion showing Clump and HP1a log2-transformed ChIP-seq enrichment and small RNA coverage in control and *clump* knockout ovaries. Only uniquely mappable reads are shown. **(C)** Scatter plots comparing normalized, 23-29 nt small RNA-seq reads mapping to TE consensus sequences in sense and antisense orientation, in *clump* KO and control ovaries. Each dot represents a TE consensus; values are average from 3 biological replicates per genotype. TEs with a Log2-normalized fold change greater than 2 are highlighted in red.

Together with the limited and variegated effects on steady-state TE RNA levels and the relatively mild phenotype of *clump* mutants, these findings suggest that Clump functions as a modulator of heterochromatin whose absence results in local perturbations; such perturbations may trigger mechanistically distinct events — such as local loss of *pogo* piRNAs or piRNA-independent transcriptional activation as for *HeT-A —* that ultimately converge to steady-state TE RNA increase. Although these effects appear partially buffered by the global piRNA response, these observations are consistent with *clump* loss primarily leading to defects at the chromatin level and are compatible with a model where Clump is mechanistically linked to HP1a-mediated transcriptional silencing.

### Clump is recruited by HP1a and acts to maintain HP1a dynamics and repressor function

To address the mechanistic role of Clump in heterochromatin, we further examined its interplay with HP1a. First, we tested how Clump and HP1a depend on each other for their localization on chromatin. It is well established that HP1a binds the conserved heterochromatin mark H3K9me2/3 via its chromodomain(Bannister et al. 2001; Lachner et al. 2001; Jacobs and Khorasanizadeh 2002). To test if Clump localization depends on HP1a, we analyzed the subcellular localization of Clump-GF upon HP1a knockdown in the germline via an HP1a shRNA activated in nurse cells by the oskar-Gal4 driver. Results showed complete loss of Clump nuclear foci and diffuse cytoplasmic signal, suggesting that HP1a is largely required for Clump localization to chromatin (Figure 4A). To determine whether HP1a can recruit Clump directly, we utilized an artificial tethering assay in which a LacI-HP1a fusion protein is targeted to a reporter containing *LacO* sites upstream of a constitutive promoter (nanos), embedded in a naive euchromatic locus(Sienski et al. 2015). We introduced the Clump-GF transgene into this system and employed ChIP-qPCR to assess Clump occupancy at the reporter locus upon HP1a tethering, alongside a positive control locus endogenously enriched in HP1 and Clump based on ChIP-seq data, and a random euchromatic locus normally devoid of Clump. As expected, without HP1a tethering, Clump ChIP-qPCR signal was only enriched at endogenous heterochromatin but not at the reporter locus or the euchromatic control region. However, in the presence of LacI-HP1a, Clump accumulates at the LacO reporter, confirming that HP1a can recruit Clump to chromatin (Figure 4B).

**Figure 4.**
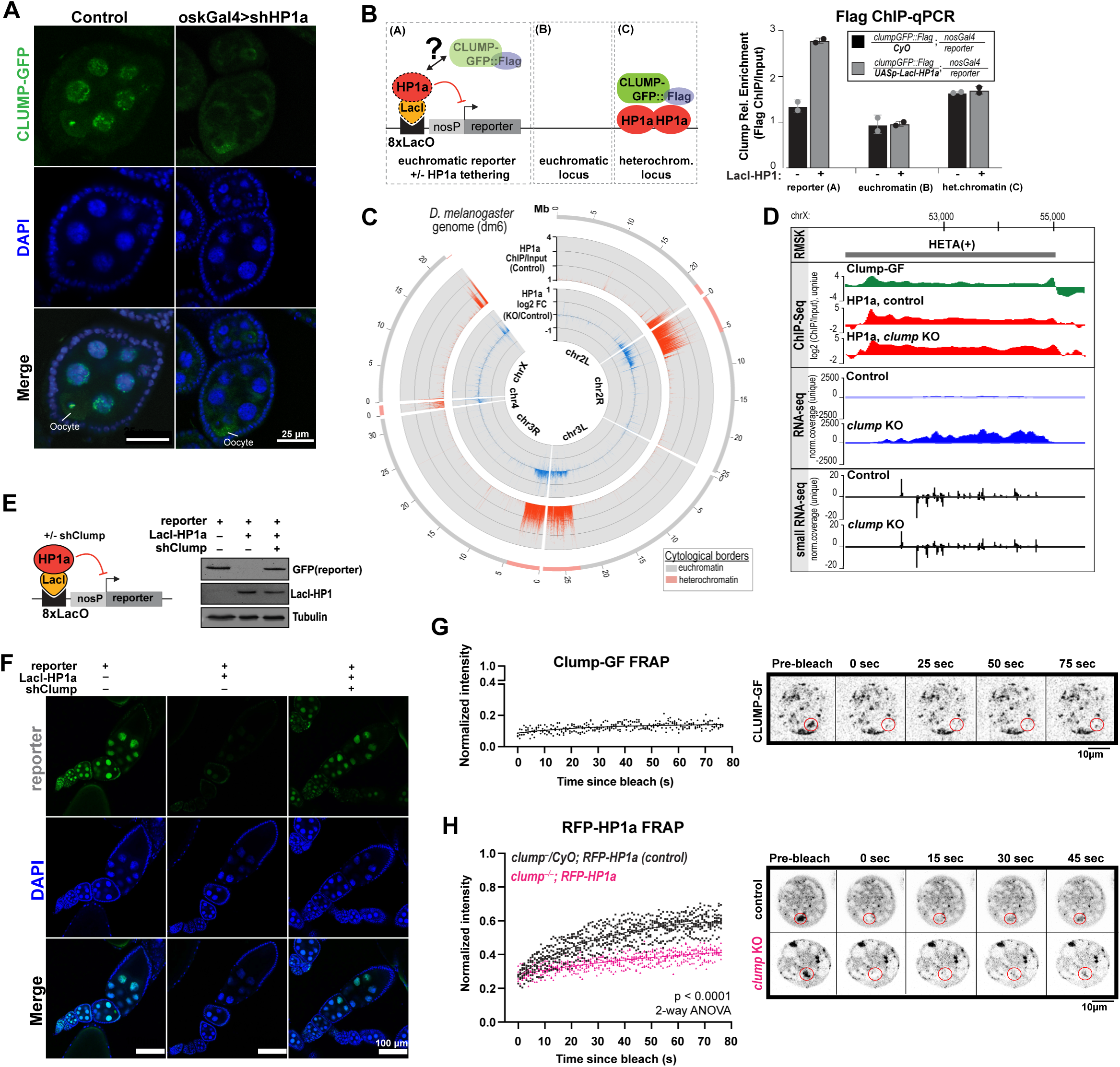
Clump contributes to HP1a silencing ability and affects HP1a mobility. **(A)** Confocal images showing Clump-GF localization in ovaries following germline knockdown of HP1a using oskar-Gal4 driver. Driver-only siblings without shHP1a expression served as controls. **(B)** ChIP-qPCR analysis for Clump-GF occupancy upon LacI-HP1a tethering to a reporter locus with LacO arrays in the promoter region. (Left) Diagram of the assay; an HP1a-devoid region (euchromatin), and endogenously HP1a– and Clump-enriched regions (heterochromatin) serve as controls. (Right) ChIP-qPCR results; error bars show s.d. from two biological replicates. **(C)** Circle plot showing HP1a enrichment across the genome in control (-/CyO) ovaries alongside with HP1a fold change in *clump* KO (-/-) vs control heterozygous (-/CyO) ovaries; n=3 biological replicates. **(D)** UCSC genome browser screenshot of a HeT-A locus with indicated RNA-seq and ChIP-seq signals in *clump* KO and control ovaries. **(E)** Schematic of the LacO/LacI-HP1a tethering reporter system and WB analysis of GFP reporter expression in indicated genotypes. **(F)** Confocal images of ovaries of indicated genotypes, as in E. **(G)** Representative images and quantification of FRAP for endogenous Clump-GF in live nurse cell nuclei. Normalized fluorescence intensity within the bleached region was measured over time after photobleaching. Representative pre-bleach and post-bleach images at the indicated time points are shown. **(H)** Representative images and quantification of FRAP for RFP-HP1a in live nurse cell nuclei. Quantification was performed on n = 9 nuclei per condition. Statistical significance was assessed by two-way ANOVA.

We next asked if Clump loss reciprocally affects HP1a chromatin distribution. Confocal imaging did not show any gross differences of HP1a subcellular distribution upon *clump* loss (Figure S4A). We further investigated potential quantitative and fine-scale effects of *clump* loss on HP1a’s genomic distribution by performing HP1a ChIP-seq in KO and heterozygous control ovaries in 3 biological replicates. Remarkably, loss of Clump leads to a slight yet consistent increase in HP1a binding across most heterochromatic regions (Figure 4C). This increase can be observed even in regions with a significant increase of TE activity, such as a subtelomeric region containing degenerate *HeT-A* copies (Figure 4D). These findings suggest that Clump is not required for HP1a’s localization but is important for its silencing function. To further test this model directly, we examined the effect of Clump depletion on the ability of HP1a to silence an artificial target, using the LacI-HP1a/LacO-reporter genetic system described above. Results showed that while LacI-HP1a efficiently silences the LacO-reporter locus in the presence of Clump, Clump knockdown reverses reporter silencing (Figure 4E,F). Taken together, these findings indicate that Clump is a heterochromatin factor that promotes HP1a’s silencing ability, effectively functioning as a corepressor.

HP1 proteins can phase separate, and heterochromatin was demonstrated to display liquid-droplet-like properties with functional implications(Larson et al. 2017; Strom et al. 2017). Considering Clump’s interplay with HP1a, we examined the mobility of Clump-GF, as well as the mobility of RFP-HP1a in *clump* KO ovaries, using Fluorescence Recovery After Photobleaching (FRAP) in live nurse cell nuclei. Clump-GF displayed minimal recovery after photobleaching, suggesting that unlike dynamic liquid droplets, Clump foci represent solid or gel-like structures with minimal molecular turnover (Figure 4G). In line with this, efforts to purify full-length Clump from *E.coli* suggested that this protein has high propensity for aggregation (data not shown). In contrast, and consistent with previous reports(Strom et al. 2017; Jankovics et al. 2018), HP1a is partially mobile, recovering to ∼60% after 75 seconds (Figure 4H). Strikingly, *clump* KO significantly reduced HP1a’s recovery rate, indicating that HP1a becomes less mobile and more stably bound to heterochromatin in the absence of Clump (Figure 4H). This reduced mobility provides a possible explanation of the higher enrichment of HP1a in *clump* KO ovaries observed by ChIP-seq. Overall, these data demonstrate that Clump is essential for maintaining the dynamic, mobile state of HP1a at heterochromatin and maintaining its repressor function.

### Clump autoregulates to restrict its intrinsic aggregation

Heterochromatin spreading is dose-dependent, as shown by position effect variegation, highlighting the need for homeostatic mechanisms that preserve epigenetic fidelity (Eissenberg et al. 1992; Allshire and Madhani 2018). Previous work highlighted Clump as a putative heterochromatin factor regulated by an H3K9me3/HP1a-dependent negative feedback mechanism, as the *clump* gene has H3K9me3 and HP1a enrichment in its promoter region and gets upregulated upon H3K9me3 depletion(Ninova et al. 2020). The finding that Clump protein itself extensively accumulates at its promoter (Figure 1J, Figure 5A) provides strong support for an autoregulatory mechanism where Clump directly regulates its own gene’s transcription. To test this explicitly, we utilized *clump* shRNA knockdown lines, which eliminate mature mRNA and protein while leaving nascent transcripts intact, thereby enabling the readout of transcriptional activity at the locus. RT-qPCR showed that loss of *clump* mRNA/protein leads to transcriptional upregulation, providing direct evidence of negative feedback (Figure 5B).

**Figure 5.**
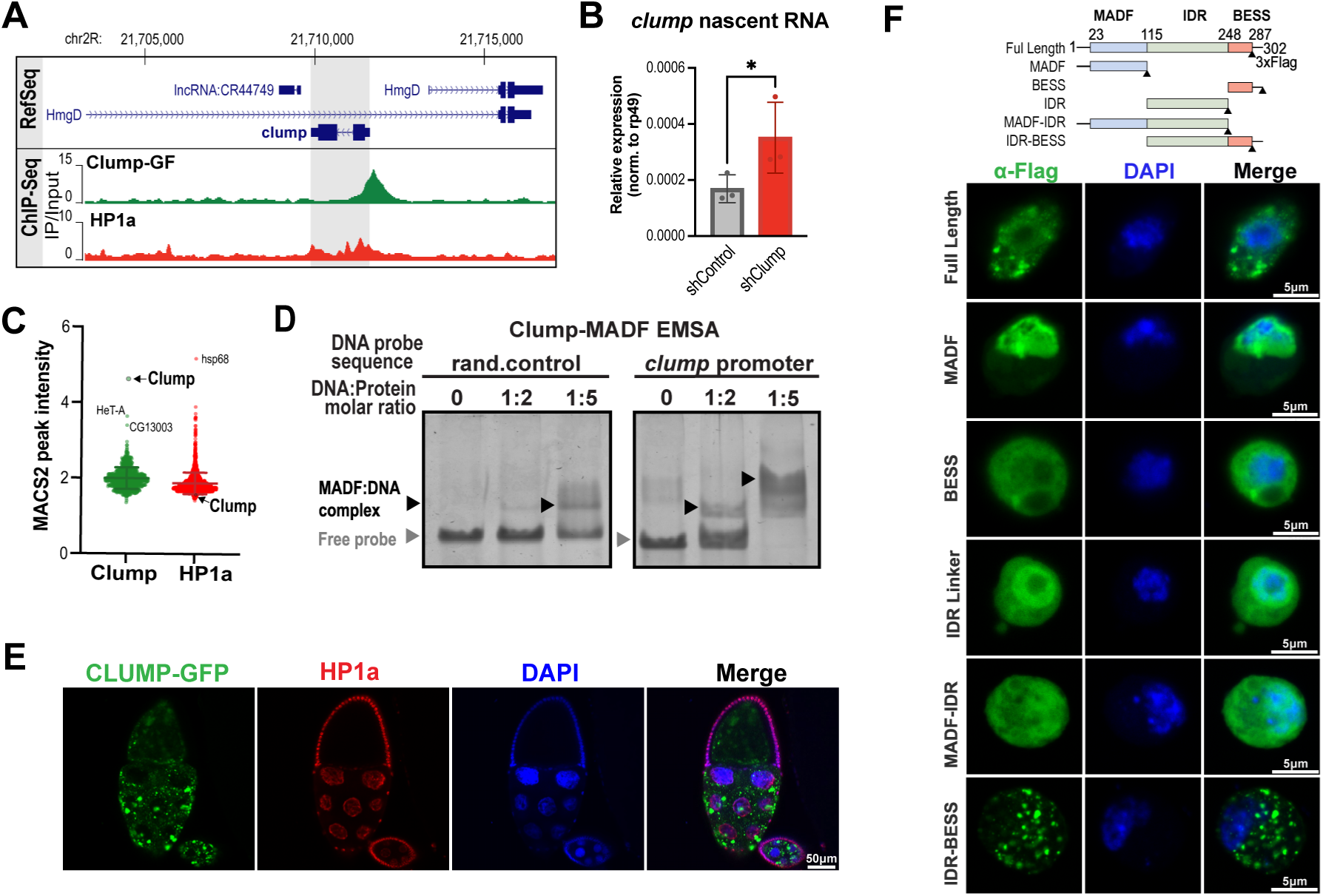
Clump autoregulates to restrict its intrinsic aggregation. **(A)** UCSC Genome Browser screenshot of the *clump* locus showing Clump-GF and HP1a ChIP-seq enrichment profiles. **(B)** RT-qPCR analysis of nascent *clump* RNA in control (shWhite) and *clump* KD (shClump) by RNAi in the germline. Error bars indicate s.d.from 3 biological replicates; *p<0.01, Student’s t-test. **(C)** ChIP-seq enrichment of Clump and HP1a at genome-wide binding sites identified by MACS2. Horizontal bars indicate the mean ± SD. **(D)** EMSA analysis showing binding of the Clump MADF domain to an oligonucleotide sequence corresponding to the *Clump* peak summit in the gene promoter region, and a random GFP-derived oligonucleotide control. **(E)** Confocal images of ovaries overexpressing GFP-Clump driven by MT-GAL4, and co-stained with anti-HP1a antibody. **(F)** Top: schematic of the full-length Clump and truncation constructs used for domain aggregation analysis. Bottom: S2 cells overexpressing the indicated FLAG-tagged Clump domains stained with FLAG antibody and AlexaFluor-488-conjugated secondary antibody.

Remarkably, the *clump* promoter exhibits an unusually high Clump:HP1a ratio compared to the rest of the genome — the Clump ChIP enrichment signal is several times higher than the genome-wide average, while the HP1a signal is lower than the genome-wide average (Figure 5C). Although HP1a appears to genetically govern the bulk of Clump localization (Figure 4A,B), this specific enrichment indicated that the *clump* locus might employ a specialized, direct mechanism for its own autoregulation. The MADF domain was shown to function as a DNA-binding module in other proteins(Cutler et al. 1998). Structural modeling of the interaction between the Clump MADF domain and dsDNA using AlphaFold3(Abramson et al. 2024) consistently yields a structure where the third α-helix is positioned in the major groove of the DNA double helix, similarly to the canonical Myb-family DNA-binding domains(Ogata et al. 1994)(Figure S5A). Following these observations, we performed an electrophoretic mobility shift assay (EMSA) using *in vitro* purified Clump MADF domain. Results confirmed that Clump MADF domain can bind DNA and indicated a preference to the sequence corresponding to the ChIP-seq summit at its own promoter over a random sequence (Figure 5D). Together, these findings uncover a molecular mechanism of Clump negative feedback autoregulation in which the protein binds its own promoter to limit its expression.

The strict autoregulatory control of *clump* transcription raises the question of why such precise dosage maintenance is required. To address this, we examined the consequences of overriding this endogenous autoregulation by ectopic overexpression, utilizing a transgenic line encoding a GFP-tagged Clump under the control of the UAS promoter, and the maternal tubulin GAL4 driver active in nurse cells. We found that Clump overexpression results in aberrant, large aggregate-like structures, which remain mostly restricted to the cytoplasm and fail to colocalize with HP1a (Figure 5E, Figure S5B). Systematic testing in S2 cells ruled out the protein tags as the source of aggregation, as aggregation was observed with 3xFLAG tag, and GFP N– and C-terminal tags (Figure 5F,S5C). To further determine the molecular basis of this behavior, we assessed the subcellular localization patterns of different truncated constructs, which revealed that Clump aggregation is driven specifically by the IDR-BESS region (Figure 5F). Notably, considering that the full-length Clump readily forms insoluble aggregates when expressed in bacteria (data not shown), this behavior likely represents a strong intrinsic biophysical property of the protein. Collectively, these data suggest that Clump is an inherently aggregation-prone protein that evolved a strict autoregulatory mechanism. By keeping its concentration below the threshold of unchecked aggregation, Clump accurately governs its own physiologically appropriate state within the nucleus to support the biophysical properties of heterochromatin. This provides a clear example of how cells utilize direct transcriptional feedback to manage the phase behavior of intrinsically aggregation-prone proteins.

## Discussion

Heterochromatin is a deeply conserved compartment of eukaryotic genomes, primarily functioning to maintain structurally condensed and transcriptionally inhibited states at repeat-rich loci(Grewal and Jia 2007; Elgin and Reuter 2013; Allshire and Madhani 2018; Grewal 2023). While HP1a is the major hallmark and effector of heterochromatin from yeast to humans, increasing evidence argues that heterochromatin is regulated non-uniformly via more nuanced, local mechanisms. In this work, we identified Clump/CG30403 as a novel, tissue-specific partner of HP1a in the *Drosophila* ovary that regulates its dynamics and silencing ability. By reinforcing HP1a-mediated TE silencing, Clump prevents TE mobilization and subsequent DNA damage, thereby acting as a safeguard for germline genome integrity and fertility. This work provides mechanistic insights that not only illuminate the specific roles of Clump but also offer a broader understanding of the dynamic regulation of heterochromatin.

We identified Clump as a novel heterochromatin-associated repressor in the *Drosophila* ovary. Using orthogonal genetic approaches, we demonstrate that Clump facilitates HP1a silencing at both artificially induced and physiological heterochromatic loci, including specific TEs, most notably *HeT-A* and *pogo*. In reproductive tissues, TE silencing is largely mediated by the piRNA pathway, which operates through positive feedback where piRNA-guided TE degradation drives further piRNA production and loading of cytoplasmic and nuclear effector PIWI proteins to in turn fuel both further post-transcriptional and H3K9me3/HP1-dependent co-transcriptional TE silencing (reviewed in (Ozata et al. 2019; Wang et al. 2023)). Reinforcing this circuit, heterochromatin itself is required for dual-stranded piRNA cluster transcription, creating a highly complex regulatory system(Rangan et al. 2011; Mohn et al. 2014). Consequently, any increase in TE steady-state RNA levels following loss of a nuclear effector may arise either from defective piRNA biogenesis and heterochromatin establishment, or from a downstream failure in heterochromatin function. Our findings support the latter model. Even though Clump loss causes selective defects in piRNA production (e.g., loss of *pogo* piRNAs), global piRNA biogenesis remains largely intact. Instead, Clump physically and genetically interacts with HP1a, placing it downstream of HP1a (see below). Further consistent with heterochromatin defect, Clump loss is associated with increased canonical transcription within piRNA clusters, and piRNA-independent upregulation of several TEs, including the telomeric *HeT-A* and *TAHRE*. Notably, telomeric TEs are the most sensitive TE to HP1a depletion(Teo et al. 2018). Together, these findings indicate that Clump is required to support effective silencing in HP1a-marked heterochromatin, rather than its establishment.

Unlike many piRNA pathway mutants, which are characterized by widespread TE upregulation and complete sterility, *clump* mutants have a relatively mild phenotype — limited increases in TE RNA levels and subfertility — despite genome-wide occupancy at TEs and heterochromatin. We propose that this reflects buffering by a largely intact piRNA pathway which masks underlying defects in HP1a-dependent silencing. In line with this, *pogo* silencing is variegated at the single-cell level, and Clump depletion results in an age– and heat stress-dependent decline in fertility without a discrete block in oogenesis or embryogenesis. Together, these findings suggest that Clump loss causes stochastic silencing failures rather than global heterochromatin collapse, with fertility defects accumulating over time. The relatively mild phenotype may also reflect partial redundancy with other MADF-BESS family members, consistent with a role for Clump in reinforcing, rather than establishing, heterochromatin-mediated silencing. More broadly, these findings highlight how heterochromatin integrity is maintained not only by core silencing machinery but also by auxiliary factors. Even though disruption of such factors may have subtle effects in laboratory-bred individuals, they can have significant biological consequences in wild populations over time.

How does Clump cooperate with HP1a? We found that Clump interacts with HP1a’s CSD via its IDR, and broadly co-localizes with it across canonical heterochromatin domains, including interspersed transposons, piRNA clusters, and a small subset of gene promoters. Genetic and artificial tethering experiments established a clear recruitment hierarchy: in the absence of HP1a, Clump fails to localize to the nucleus and diffuses into the cytoplasm, whereas tethering HP1a to euchromatin is sufficient to ectopically recruit Clump. Conversely, Clump loss does not grossly alter HP1a’s genomic occupancy. However, Clump loss significantly reduces HP1a mobility. HP1a is well known to undergo phase separation, driving the condensation of heterochromatin into membraneless domains with liquid-like properties(Larson et al. 2017; Strom et al. 2017). Clump is highly disordered and interacts with HP1a via an IDR, which represents a structural motif frequently employed to fine-tune multivalent interactions within condensates. Thus, our data suggest that Clump may act as a biophysical modulator of heterochromatic domains. Interestingly, Clump itself appears relatively immobile, suggesting it may play a scaffolding role. Ultimately, the mechanistic link between the physical state of heterochromatin and its biological function remains poorly understood. Clump offers a unique model to explore how biophysical properties influence silencing in the future.

Finally, we identified a remarkable autoregulatory mechanism governing Clump’s expression. Clump is enriched at its own promoter and inhibits its transcription, consistent with a negative feedback loop and suggesting that Clump is a dosage-sensitive factor. We observed that Clump overexpression leads to severe cytoplasmic aggregation, which may underlie the necessity for its levels to be tightly constrained. Together, these findings suggest that autoregulation serves to maintain Clump within a narrow physiological range compatible with proper phase behavior and heterochromatin function, highlighting how precise control of condensate-associated factors may be critical for epigenetic stability.

## Materials and Methods

### Fly stocks and maintenance

Flies were maintained at 25°C on standard molasses media (Caltech recipe). Females were put on yeast for 2 to 3 days before ovary dissection. The following stocks were obtained from the Bloomington Drosophila Stock Center (BDSC): Nanos-GAL4-VP16 (BDSC 4937), matalpha4-GAL-VP16 (BDSC 7063), osk-GAL4-VP16 (BDSC 44242), MTD-GAL4-VP16 (BDSC 31777), RFP-HP1a (BDSC 30562), shHP1a (BDSC 36792), shWhite (BDSC 33623). The tethering line pUASp>LacI_Su(var)205 [attP40]/CyO; lacO_nos>GFP-Piwi[attP2]/TM3 was obtained from the Vienna Drosophila Resource Center (VDRC 313409)(Sienski et al. 2015). The HA-HipHop fly line(Saint-Leandre et al. 2020), was generously provided by Dr. Mia Levine.

To generate the *clump[Δ::dsRed]* KO line, we employed a CRISPR-Cas9-mediated homology-directed repair strategy(Gratz et al. 2014). Two guide RNAs (Clump-sgRNA1: GGAGCAGCAGTTGCTCGAAG and Clump-sgRNA2: GCTCCATAGAGAGCATATAC) were designed using the online CRISPR design tool at: http://targetfinder.flycrispr.neuro.brown.edu/ to generate double-strand breaks near the transcription start site and downstream of the termination site of *clump* gene. These two sgRNAs were cloned into the pCFD5 plasmid based as described previously (Port and Bullock 2016). The donor construct was generated using pHD-DsRed (Addgene #51434) as the backbone(Gratz et al. 2014); the entire *clump* gene was deleted and replaced with 3xP3-dsRed cassette, flanked by ∼1kb upstream (left donor) and downstream (right donor) homology arms. To generate the *clump[KI.eGFP::3xFLAG]* (Clump-GF) knock-in line, we used the same procedure and sgRNAs, except the donor plasmid contained an eGFP-3xFLAG tag inserted before the stop codon, with the 3XP3-DsRed marker placed downstream of the *clump* gene for selection. For all CRISPR lines, donor and sgRNA plasmids were injected into yw;;nos-Cas9(III-attP2) embryos by BestGene Inc., where G0 transformants were back crossed to yw. To establish a stable, homozygous transgenic KO line, dsRed-positive male was isolated and crossed to a standard CyO balancer stock. The resulting dsRed/CyO progeny were then intercrossed. Correct CRISPR integrations were confirmed by long-read sequencing of target PCR amplicons. Wild type controls are sister yw lines.

To generate the pUASp-shClump lines, target sites were selected using the Designer of Small Interfering RNA tool (http://biodev.extra.cea.fr/DSIR/DSIR.html)(Vert et al. 2006). Two shRNA sequences, AACATCAATTCCTCAACTAAA and TTTAGTTAGAAATATGTCCAT, were cloned into the pVALIUM20 plasmid(Ni et al. 2011). For the pUAS-Clump overexpression lines, GFP-Clump fusion constructs were introduced into the pUASP vector pUASP (DGRC Stock 1189; https://dgrc.bio.indiana.edu//stock/1189; RRID:DGRC_1189) via Gibson Assebmly. The resulting plasmids were injected into the *y[1]w[67c23];P{y[+t7.7]=CaryP}attP2* strain (BDSC 8622) or the *y[1]w[67c23]; P{CaryP}attP40* for phiC31 transformation at BestGene Inc. Correct integrations were verified by sequencing of target PCR amplicons.

### S2 cells expression plasmids

Expression vectors encoding GFP– or FLAG-tagged fusion proteins of different Clump and HP1a fragments were generated by cloning indicated fragments in a custom vector containing the ubiquitin promoter derived from Addgene plasmid #74280 using NEBuilder® HiFi DNA Assembly.

### S2 cells maintenance and transfection

S2 cells were cultured at 25°C in Schneider’s *Drosophila* Medium supplemented with 10% heat-inactivated FBS and 1x Penicillin-Streptomycin. To express tagged proteins, indicated constructs were transfected by TransIT®-Insect Transfection Reagent (Mirus, 6105) according to the manufacturer’s instructions. Cells were harvested 48-72hr post-transfection for downstream analysis.

### RNA preparation, qRT-PCR and data analysis

Total RNA was isolated from ovaries collected at specific ages using TRIzol (Invitrogen). 1 µg RNA was treated with DNAseI (Invitrogen), and reverse transcribed using the LunaScript RT SuperMix Kit (Primer free). OligodT primer was used for RT–qPCR analysis of *Clump* knockdown efficiency and transposon expression. Random hexamer primer was used to detect spliced and unspliced transcripts from the 42AB and Sox102F piRNA clusters and nascent *Clump* transcripts. Primer sequences are listed in Table S1. Primers used to detect spliced and unspliced piRNA cluster transcripts were adapted from Zhang et al(Zhang et al. 2014), qPCR was performed using PowerTrack™ SYBR Green Master Mix from Applied Biosystems™. The qPCR primers are listed in Table S1. Experiments were performed in 3 independent biological replicates, and CT values were calculated from technical duplicates or triplicates, *rp49* gene was used to serve as an internal control. Statistical significance was calculated by two-tailed Student’s t-test.

### RNA-seq library preparation and sequencing

2-4 days adult *clump[Δ::dsRed]* and *clump[Δ::dsRed]/CyO* siblings were collected and fed yeast for 2 days before ovary dissection. RNA was isolated from samples of 100 pairs of ovaries in 3 biological replicates per genotype using TRIzol (Invitrogen). The library preparation was done by UCI Genomics Research and Technology Hub with PolyA selection by using Illumina TruSeq Stranded mRNA Kit (Illumina: 20020594). The libraries were sequenced on Illumina NovaSeq 6000 at the UCI Genomics High Throughput Facility with 30 million 100-bp paired-end reads.

### Small RNA-seq library preparation and sequencing

2-4 days adult females were collected and fed yeast for 2 days before ovary dissection. RNA was isolated from samples of 100 pairs of ovaries in 3 biological replicates per genotype using TRIzol (Invitrogen). Small RNA-seq library construction was carried out after rRNA depletion using the NEBNext Small RNA Library Prep Set for Illumina (E7330S) following manufacture’s protocol. Libraries were sequenced on the Illumina NovaSeq 6000 at the UCI Genomics High Throughput Facility with approximately 30 million 150-bp paired-end reads. Because the informative small RNA sequence was contained in Read 1, only Read 1 was used for downstream analysis.

### ChIP-seq and ChIP-qPCR

2-4 days adult flies of desired genotypes were collected and fed yeast for 2 days before dissection. A total of 200 pairs of ovaries per sample, in two or three biological replicates, were dissected and placed in ice-cold PBS and fixed in 1% formaldehyde for 10 min. Fixation was quenched with 125mM glycine, and samples were washed twice with 1ml PBS. Samples were then washed twice in 1ml Farnham buffer (5mM Hepes PH 8.0, 85mM KCl, 0.5% NP40) with 10 strokes of a plastic pestle. After centrifugation, the pellet was resuspended in 1 mL RIPA (standard recipe) and sonicated on ice using a Branson sonicator (8 cycles of 15s on / 60s off at 4°C). Sonicated lysate was centrifuged at 13,300 rpm for 20min at 4°C. Chromatin immunoprecipitation was performed using the following antibodies: anti-HP1a (DSHB C1A9), anti-FLAG (Sigma-Aldrich F3165), anti–GFP (Invitrogen A-11122). For each sample, antibodies were incubated with 50ul Dynabeads Protein G overnight at 4°C with rotation. Beads were washed five times for 10min each with a wash buffer (10mM Tris PH 7.5, 500mM LiCl, 1% NP40, 1% Sodium deoxycholate). To reverse crosslink, 200 uL proteinase K buffer (200mM Tris 7.4, 25mM EDTA, 300mM NaCl, 2% SDS, 0.5mg/ul proteinase K) was added, followed by incubation at 55°C for 2h and 65°C overnight. DNA was purified by phenol-chloroform extraction and quantified using a Qubit fluorometer. DNA quality and size distribution were assessed using High Sensitivity DNA ScreenTape on an Agilent 4150 TapeStation System.

ChIP-seq library construction was carried out using the NEBNext Ultra™ II DNA Library Prep Kit for Illumina (E7645). Libraries were sequenced on the Illumina NovaSeq 6000 at the UCI Genomics High Throughput Facility with approximately 30 million 150-bp paired-end reads. ChIP-qPCR was performed on a CFX Connect Real-Time PCR Detection System. For ChIP-qPCR analysis, Cq values were normalized to *rp49* internal control, and further normalized as ChIP to Input. Relative enrichment was calculated using the ΔΔCq method.

### Tissue dissection and Western Blot analysis

Different tissues and different stages of embryos were collected and lysed with RIPA buffer (20mM Tris PH7.4, 150mM NaCl, 1% NP40, 0.5% Sodium deoxycholate, 0.1% SDS) supplied with protease inhibitor on ice for 20 min. The samples were then centrifuged to remove debris. Protein amount was quantified by BCA assay (Thermo, 23225). Equal amounts of protein were loaded for Western Blot analysis. The following antibodies and concentrations were used: anti-FLAG (Sigma F3165, 1:5000), anti-Tubulin (Sigma T5168, 1:10000), anti-HP1a (Developmental Studies Hybridoma Bank C1A9, 1:1000), anti-GFP (Gift from Alexei Aravin, 1:5000), Anti-mouse IgG (Cell signaling 7076, 1:2000), Anti-Rabbit IgG (Cell signaling 7074, 1:2000).

### Co-immunoprecipitation

For ovarian lysates, 2-4 days adult flies were collected and fed yeast for 2 days before dissection. 50 pairs of dissected ovaries per sample were lysed on ice using a dounce homogenizer in 1ml lysis buffer (20mM Tris PH7.4, 150mM NaCl, 10% glycerol, 0.4% NP-40, 0.1% triton-X) supplied with protease inhibitor (Roche, 11697498001). For S2 cells, cells were harvested after 48-72h transfection, washed with PBS, and lysed with lysis buffer. For HP1a immunoprecipitation to assess the interaction between Clump-GFP and HP1a, lysates were incubated with 10 µL anti-HP1a (Developmental Studies Hybridoma Bank C1A9) overnight, followed by incubation with PBS-washed Dynabeads™ Protein G (Invitrogen, 10003D) for 2h. Beads were washed four times with a wash buffer (20mM tris PH 7.4, 150mM NaCl, 0.4% NP-40) and eluted by boiling in a 45 µl 1X LDS sample buffer. The supernatant was used for Western Blot analysis.

### Confocal Imaging of fixed ovaries and cells

Ovaries of flies expressing fluorescently tagged proteins were dissected at specified ages following 2 days of yeast feeding. Samples were fixed in 4% formaldehyde for 20 min at room temperature, washed three times for 5 min each in PBS, and mounted in VECTASHIELD® Antifade Mounting Medium (H-1000) with DAPI. To stain S2 cells, cells were fixed in 4% formaldehyde for 20 min at room temperature and washed three times with PBS-TX buffer (PBS, 0.1% Tween-20, 0.3% Triton X-100). Cells were blocked in 4% BSA in PBS-TX for 1 h at room temperature, then incubated overnight at 4°C with anti-FLAG antibody (Sigma-Aldrich F3165, 1:1000) diluted in blocking buffer with rotation. Cells were washed three times for 5 min each in PBS-TX and incubated with Alexa Fluor 488-conjugated anti-mouse secondary antibody (Invitrogen A-11001, 1:400) for 1.5 h at room temperature. After three additional washes with PBS-TX, cells were resuspended in 30 µl VECTASHIELD mounting medium and imaged to assess protein localization. Imaging was performed on a Zeiss LSM 880 inverted confocal microscope. Images were processed in Fiji(Schindelin et al. 2012).

### Bioinformatic analysis

The *D.melanogaster* genome assembly dm6 was used for all analysis. For RNA-seq differential expression analysis, a combined kallisto index was generated from dm6 annotated transcripts and RepBase consensus transposon sequences. Reads were then pseudoaligned to this combined reference to quantify both gene and transposon expression (Bao et al. 2015). Differential expression analysis was performed using sleuth with default settings(Pimentel et al. 2017). Normalized genome coverage tracks (fragments per million mapped reads) were generated on uniquely mapping read pairs using bamCoverage from deepTools (Ramirez et al. 2016).

For ChIP-seq analysis, reads were aligned to dm6 with STAR with the parameters ––alignIntronMax 1 ––alignEndsType EndToEnd ––outFilterMultimapNmax 1 ––outSAMtype BAM SortedByCoordinate. Peak analyses were performed using MACS2 (Zhang et al. 2008), and peaks annotation was performed using ChIPseeker (Yu et al. 2015). For genome-wide Clump-GF enrichment visualization by circle plots, the dm6 genome was partitioned into 5-kb intervals. ChIP/Input signal was defined as the ratio of RPM-normalized ChIP to Input values (ChIP/Input) in each 5kb interval. Heterochromatin boundaries were obtained from Riddle et al. and converted to dm6 coordinates using the UCSC Genome Browser LiftOver tool (Riddle et al. 2011). The circular plots were generated using the Circos package(Krzywinski et al. 2009). Heatmaps were generated by deepTools(Ramirez et al. 2016). Normalized genome coverage tracks were generated using bamCompare from deepTools with the parameter ––operation log2. To define “euchromatic TE loci”, RepeatMasker annotations were filtered against cytological heterochromatin boundaries and piRNA clusters.

For small RNA-seq analysis, adapters were removed by cutadapt(Martin 2011). Then reads were aligned to rRNA first to filter rRNA reads, and the remaining reads were aligned to dm6 genome using STAR with the parameters ––alignEndsType EndToEnd –– outFilterMultimapNmax 1 ––alignIntronMax 1 ––outSAMtype BAM SortedByCoordinate. Reads mapping to piRNA clusters were quantified using featureCounts and normalized to total reads to obtain reads per kilobase million (RPKM). To quantify transposon-derived small RNAs, reads were uniquely aligned to consensus transposon sequences (RepBase) using STAR with the parameters ––alignEndsType EndToEnd –– outFilterMultimapNmax 1 ––alignIntronMax 1. Read numbers were summarized by featureCounts and further normalized to total reads to obtain RPM values. To calculate total reads length distribution, reads with removed adapter length were calculated and normalized into RPM. The TE-mapped reads length distribution and ping-pong signatures were analyzed using piPipes with default parameters(Han et al. 2015).

### Fertility assay

To test female fertility, five 0-2 days virgin females per replicate of desired genotypes were mated with five 2-5 days wild type males. At defined ages, flies were transferred to fresh vials every 24h. All vials were kept for 12 days before progeny were counted. Each experiment was performed with three biological replicates.

To measure egg-laying rates and embryo hatchability, five 0-2 days virgin females per replicate were mated with five 2-5 days wild type siblings in small cages for 24 h. Embryos were collected on fresh apple-juice agar plates for 24h and then incubated for an additional 24 h. Hatch rate was calculated as the proportion of hatched larvae relative to total eggs laid. Each assay was performed in 3 biological replicates.

### Protein expression and purification of the Clump MADF domain

The sequence of Clump MADF domain was cloned into a VP13 vector containing a N-terminal His_6_-MBP tag followed by a TEV cleavage site for expression in E. coli BL21 strains. Cells were grown to an optical density at 600nm (OD600) 0.6-0.8 at 37 °C and induced with 0.1 mM isopropyl β-d-1-thiogalactopyranoside (IPTG) at 18°C for 18-24 h. The bacteria was collected and resuspend in lysis buffer (20 mM Tris, pH 7.5, 300 mM NaCl, 10 mM imidazole and 10 % (v:v) glycerol) with 10 mM phenylmethylsulfonyl fluoride (PMSF), followed by purification using a Ni^2+^-NTA column. The His_6_-MBP tag was then removed by TEV protease cleavage, and the cleaved Clump MADF was passed through a second Ni^2+^-NTA column again to remove the His_6_-MBP tag. The purified Clump MADF protein was concentrated and stored at −80 °C before use.

### Electrophoretic mobility shift assay (EMSA)

Electrophoretic mobility shift assay (EMSA) was performed following a previously published protocol (Chen et al. 2024). The control oligonucleotide probe sequence was a fragment from GFP (upper strand, 5’– ATGGTGAGCAAGGGCGAGGAGCTGTTCACCGGGGTGGTGCCCATCCTG-GTCGAGCTGGAC-3’, and the Clump promoter oligonucleotide sequence corresponds to the ChIP-seq peak region (upper strand, 5’-GGTGGAGAGAAGCAGCAGAAAAGCAGAAAGCAAC-ATGTACACCGTCAGTT-3’).

Forward and reverse oligonucleotides were first annealed in buffer containing Tris pH 7.5, 50 mM NaCl, and 5% Glycerol to generate double-stranded DNA probes. The double-stranded DNA probes and Clump MADF domain were incubated for 20 min at 4°C in a binding buffer (Tris pH 7.5, 50 mM NaCl, 5% Glycerol, 0.05% β-ME) and run on a native 8% polyacrylamide gel. The electrophoresis was performed using 1XTris-borate-EDTA (TBE) buffer at a constant voltage at 100 V for 30 min at 4°C. DNA was detected by staining with SYBR Gold and visualized by gel imager.

### Structure predictions

Protein structure predictions were performed using the AlphaFold3 web server (DeepMind) with default inference parameters and stochastic seed initialization. Protein sequences were retrieved from the UniProt database. Five independent structural models were generated for each sequence using a random seed and inspected for structural convergence. Predicted models were analyzed and visualized using UCSF ChimeraX(Meng et al. 2023).

### RNA In situ HCR

Ovaries from 2-4 day flies were dissected in PBS and fixed in 4% formaldehyde in PBS for 20min, then downstream procedures were done using the HCR™ Gold RNA-FISH kit (Molecular Instruments) according to the manufacturer’s instructions. Buffers, probes, and amplifiers for HCR v3 were purchased from Molecular Instruments. HCR probes correspond to positions 2200-3400 of *Het-A* and 100-792 of *pogo* RepBase consensuses.

### Fluorescence recovery after photobleaching (FRAP)

FRAP experiments were performed on a Zeiss LSM 880 inverted confocal microscope. An approximately 1 μm-radius region of interest was photobleached. RFP-HP1a photobleaching was done with a 594nm laser at 100% power, and GFP-Clump photobleaching was done with a 488nm laser at 100% power. Recovery images were acquired with a 0.5s interval for 90s after bleaching. Fluorescent intensity within the bleached region was quantified over time using the Zeiss LSM 880 acquisition software. For each FRAP trace, the average intensity of the first five pre-bleach frames was used as the baseline and set to 1, fluorescence recovery after bleaching was then normalized to this baseline and plotted as relative fluorescence intensity over time. Recovery curves were then plotted and analyzed in GraphPad Prism.

## Competing Interest Statement

The authors declare no competing interests.

## Acknowledgements

We are grateful to Dr Sihem Cheloufi, Dr Seàn O’Leary, and members of the Ninova Lab for scientific discussions and feedback, and UC Riverside undergraduate students Priya Kumar, Anastasiya Airapetian, Matea Ibrahim, and Hannah Holmes for technical assistance in the laboratory. We thank Mia Levine for kindly sharing the HipHop-HA transgenic fly strain, and the Aravin and Fejes-Toth labs for sharing multiple GAL4 driver strains.

## Author contributions

**M.N.**: Conceptualization, Resources, Formal analysis, Supervision, Funding acquisition, Investigation, Visualization, Methodology, Project administration, Writing — original draft, Writing — review & editing. **K.W.:** Investigation, Formal analysis, Data curation, Validation, Visualization, Methodology, Writing — original draft, Writing — review & editing. **R.C**, **A.G.:** Investigation. **J.F., J.S.:** Resources.

## Funding Sources

This study was funded by NIH grants R00HD099316 and R35GM151016 to M.N.

## Supplementary Figure Legends

**Supplementary Figure 1.**
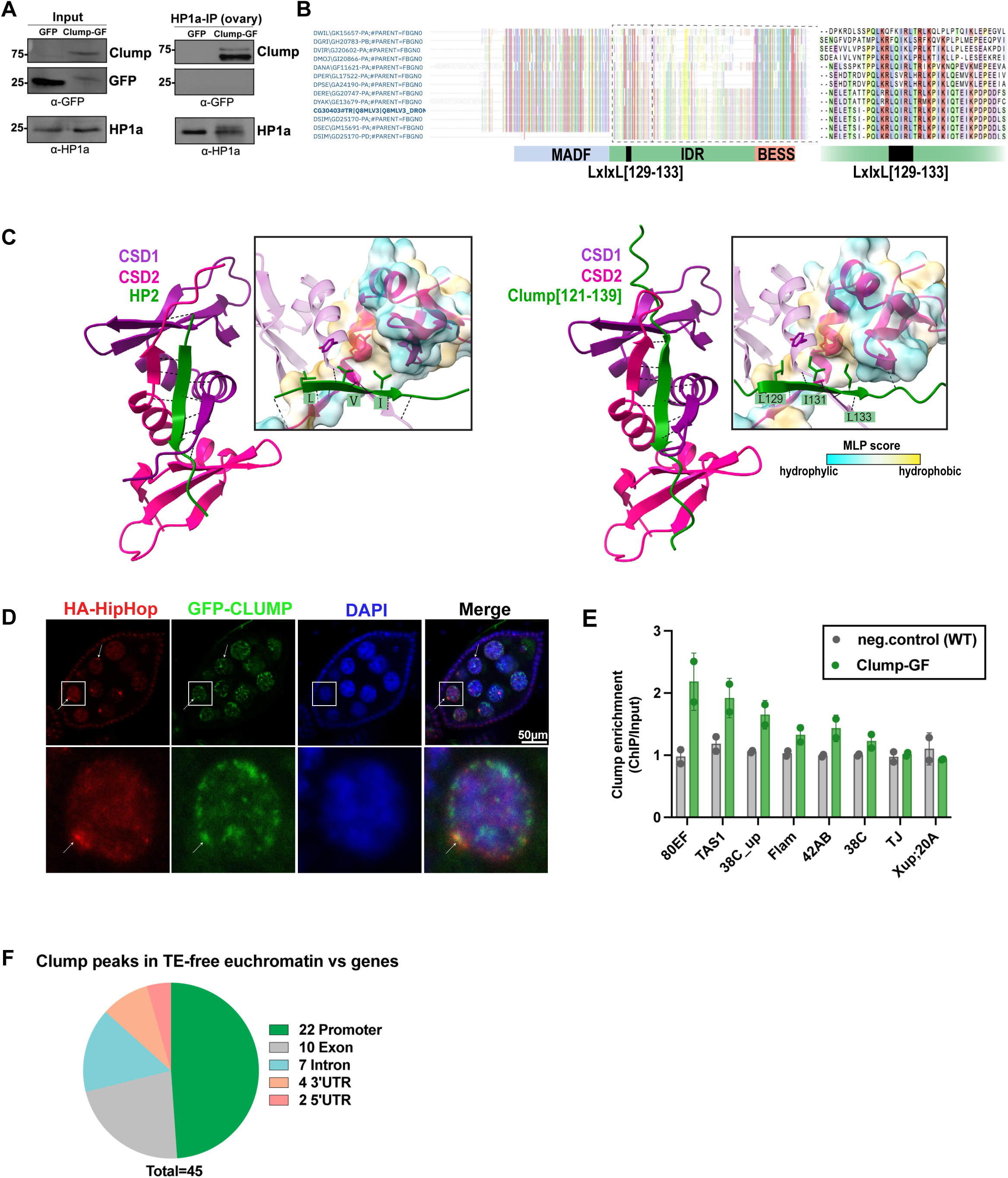
(**A**) Co-immunoprecipitation followed by WB analysis showing association of Clump-GF with HP1a in *Drosophila* ovaries. **(B)** Multiple sequence alignment of Clump homologs from 12 Drosophilid species with sequenced genomes. Amino acid residues are colored according to the clustal2 coloring convention. The position of the LxIxL motif, predicted to interact with HP1a CSD groove is indicated. **(C)** Structural models of the HP1a CSD dimer in complex with either the Clump IDR or the well-characterized HP1 CSD interactor, HP2^44,72^ for comparison. Black dashed lines denote predicted intermolecular hydrogen bonds by ChimeraX using default parameters. In expanded views, selected interacting residues are displayed as sticks, while the remaining protein structure is shown in cartoon representation; the Molecular Lipophilicity Potential (MLP) mapping on the surface of one HP1a CSD monomer is shown to illustrate the hydrophobicity of the binding pocket. **(D)** Confocal image of HA-Hiphop and GFP-Clump colocalization in stage 7 cyst. **(E)** Clump enrichment at the indicated piRNA clusters (from GFP ChIP-seq data). Wild-type ovaries were used as a negative control. **(F)** Pie chart showing the genomic annotation of 45 Clump ChIP-seq peaks located in TE-free euchromatic regions, as classified by ChIPseeker. Peaks were assigned to promoter, exon, intron, 3′ UTR, or 5′ UTR regions. The number of peaks in each category is indicated.

**Supplementary Figure 2.**
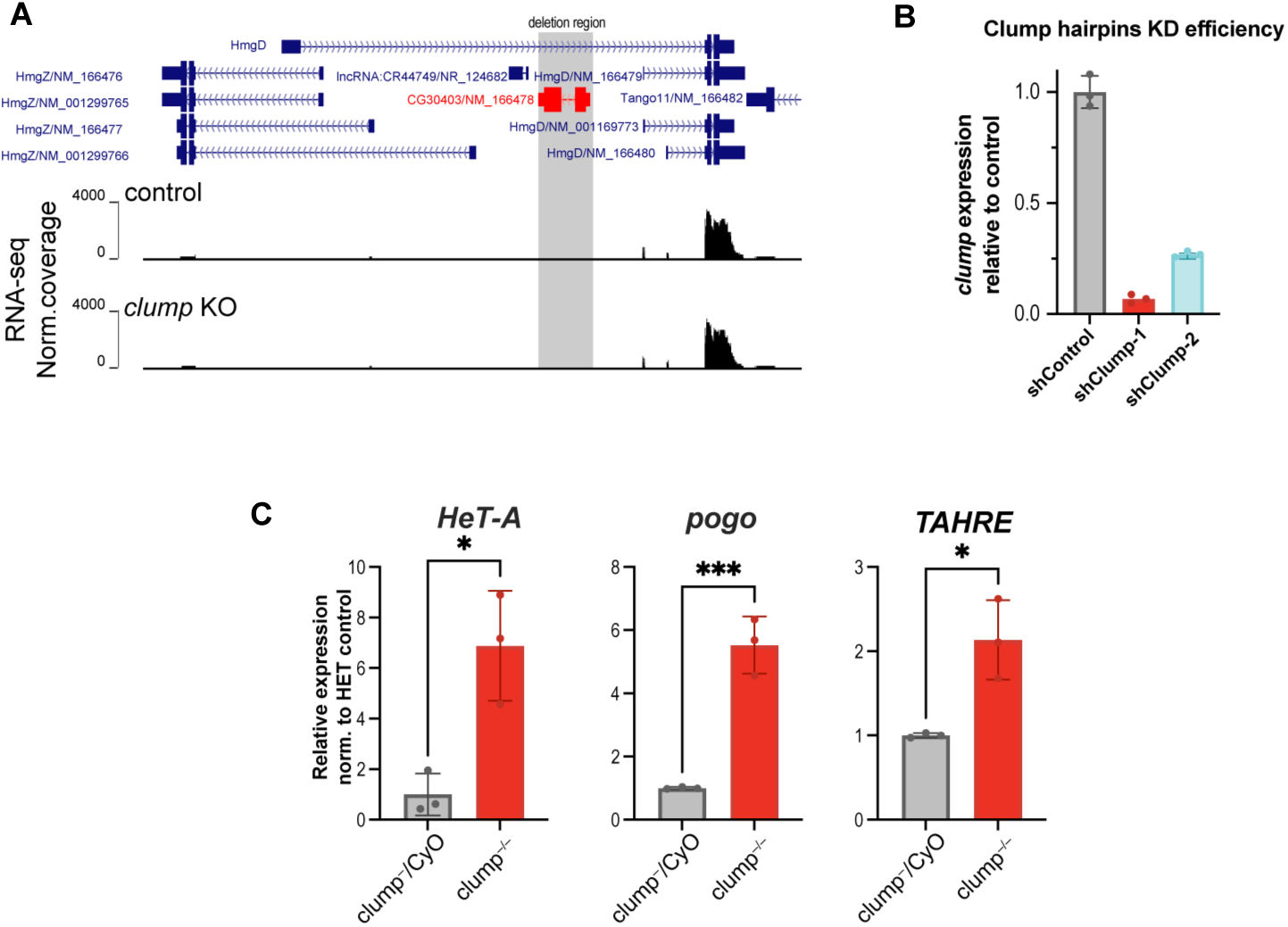
(**A**) UCSC Genome Browser view of the *clump/CG30403* locus and neighboring gene annotations. RefSeq annotations place *clump/CG30403* within an intron of a long annotated *hmgD* isoform that is not expressed. Other nearby *hmgD* isoforms expression is not affected in *clump* KO **(B)** Clump germline knockdown using two different shRNAs, shClump-1 and shClump-2, driven by nos-Gal4, reduces clump transcript levels. Individual points represent biological replicates, and error bars represent mean and SD. *: p<0.05, **: p<0.01, ***: p<0.001(Student’s t-test). **(C)** RT-qPCR analysis to validate the TE upregulation upon *clump* KO detected in RNA-seq. Individual points represent biological replicates, and error bars represent mean and SD. *: p<0.05, **: p<0.01, ***: p<0.001(Student’s t-test).

**Supplementary Figure 3.**
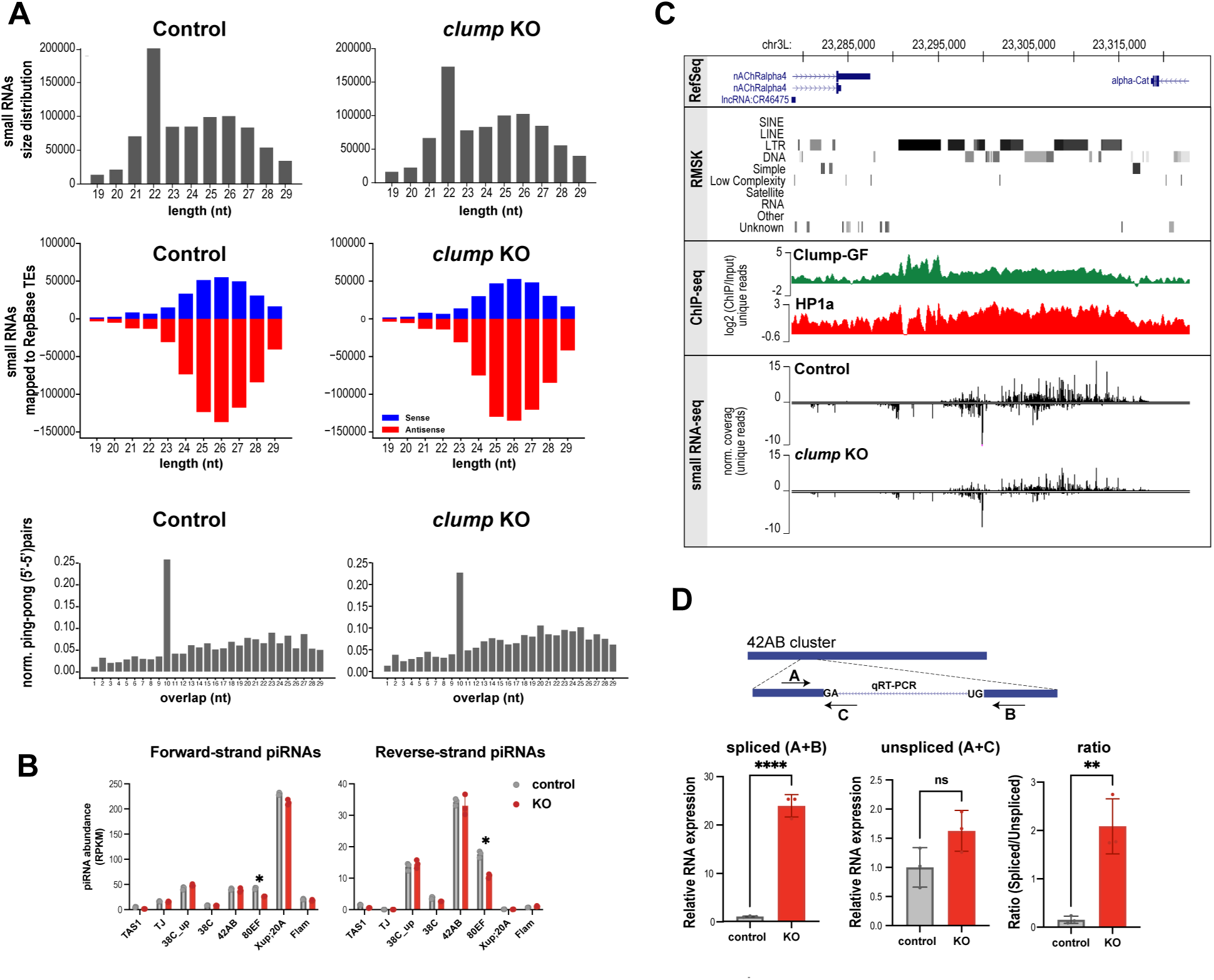
(**A**) Size distribution, strand bias, and ping-pong signature of small RNAs in control and *clump* ovaries. **(B)** Abundance of piRNAs (RPKM) mapped to the indicated piRNA clusters in control and *clump* KO ovaries. **(C)** Genome browser view of piRNA reads mapping to the 80EF cluster in control and Clump KO ovaries. **(D)** RT-qPCR analysis of spliced and unspliced transcripts from the 42AB piRNA cluster in control and *clump* KO ovaries. ns, not significant; *: p<0.05, **: p<0.01, ***: p<0.001(Student’s t-test).

**Supplementary Figure 4.**
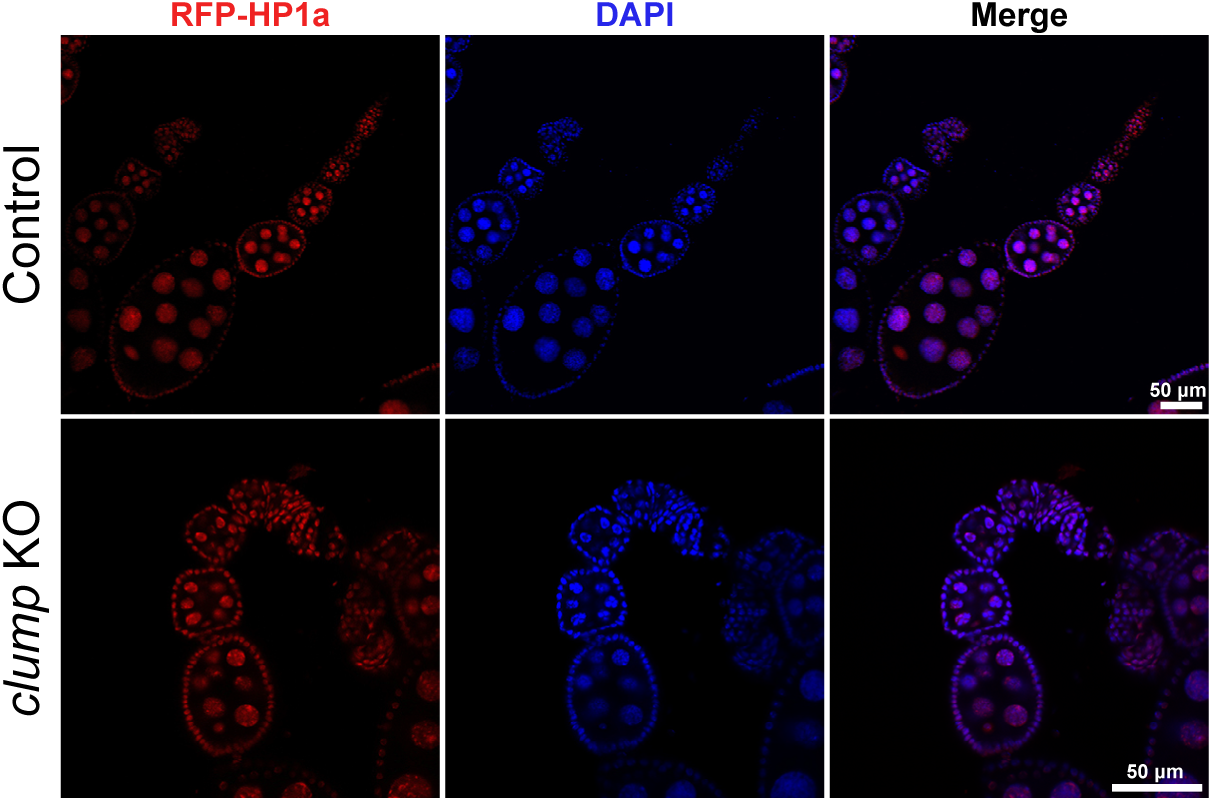
Confocal images of RFP-HP1a in control and *clump* knockout ovaries.

**Supplementary Figure 5.**
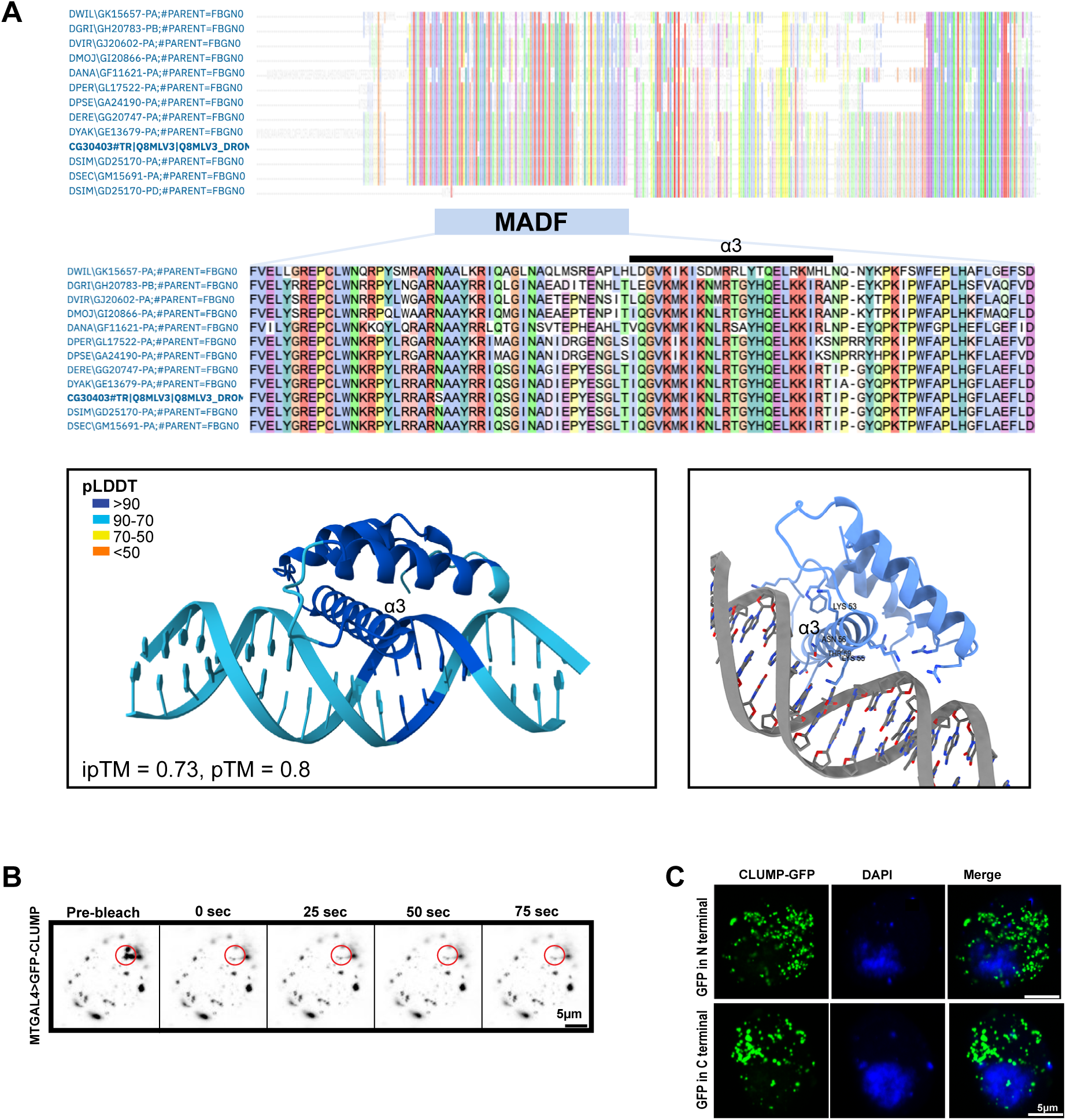
(**A**) Analysis of MADF domain and its dsDNA binding potential. (Top) Multiple sequence alignment of Clump homologs from 12 Drosophilid species with sequenced genomes with extended view on the MADF domain. Amino acid residues are colored according to the clustal2 coloring convention. (Bottom) AlphaFold3 prediction of the MADF domain bound to dsDNA. The left panel is colored by pLDDT score (confidence). The right panel shows a ChimeraX render highlighting key residues located <4Å from the DNA backbone, represented as sticks. α3 marks the third α–helix **(B)** Representative FRAP images of GFP-Clump aggregates in live fly ovaries ectopically overexpressing GFP-Clump by maternal tubulin-Gal4 (MT-Gal4) driver. **(C)** S2 cells overexpressing GFP-tagged Clump with GFP fused to either the N terminus or the C terminus.

## Notes

### Competing Interest Statement

The authors have declared no competing interest.

